# A Cytosolic Reductase Pathway is Required for Complete N-Glycosylation of an STT3B-Dependent Acceptor Site

**DOI:** 10.1101/2021.08.31.458373

**Authors:** Marcel van Lith, Marie Anne Pringle, Bethany Fleming, Giorgia Gaeta, Jisu Im, Reid Gilmore, Neil J Bulleid

## Abstract

N-linked glycosylation of proteins entering the secretory pathway is an essential post-translational modification required for protein stability and function. Previously, it has been shown that there is a temporal relationship between protein folding and glycosylation, which influences the occupancy of specific glycosylation sites. Here we use an *in vitro* translation system that reproduces the initial stages of secretory protein translocation, folding and glycosylation under defined redox conditions. We found that the efficiency of glycosylation of hemopexin was dependent upon a robust NADPH-dependent cytosolic reductive pathway, which could also be mimicked by the addition of a membrane impermeable reducing agent. The identified hypoglycosylated acceptor site is adjacent to a cysteine involved in a short range disulfide bond, which has been shown to be dependent on the STT3B-containing oligosaccharyl transferase. We also show that efficient glycosylation at this site is dependent on the STT3A-containing oligosaccharide transferase. Our results provide further insight into the important role of the ER redox conditions in glycosylation site occupancy and demonstrate a link between redox conditions in the cytosol and glycosylation efficiency.

## Introduction

Proteins entering the secretory pathway are subject to a variety of modifications, the most prevalent of which include N-linked glycosylation and disulfide formation (Bulleid, 2012; Cherepanova et al., 2016). N-glycosylation is catalysed by one of two oligosaccharyl transferases (OST) that transfer a pre-formed oligosaccharide from a dolichol-phosphate intermediate to asparagine residues on the polypeptide chain within the consensus sequence -N-X-S/T where X is any amino acid other than proline (Kelleher et al., 2003). The two OST isoforms are multi-subunit complexes characterised by the catalytic subunits STT3A or STT3B with common subunits as well as complex-specific subunits including DC2 and KCP2 for the STT3A, and the thioredoxin-domain containing proteins MagT1 or TUSC3 for the STT3B complex (Blomen et al., 2015; Roboti and High, 2012; Shibatani et al., 2005). It has been demonstrated previously that the STT3A complex associates with the ER translocon (Braunger et al., 2018; Shibatani et al., 2005) and catalyses the co-translational glycosylation of proteins, whereas the STT3B complex glycosylates sites skipped by STT3A acting predominantly post-translationally (Cherepanova et al., 2014; Ruiz-Canada et al., 2009). Because of their distinct specificities, some substrates require the STT3A or STT3B complexes for efficient glycosylation (Cherepanova and Gilmore, 2016; Cherepanova et al., 2014). Indeed, recent proteomic analysis of glycoproteins synthesised in either STT3A or STT3B depleted cells identify classes of STT3A and STT3B dependent N-glycosylation sites (Cherepanova et al., 2019). Deficiency of the STT3B complex cannot be compensated by the STT3A complex, resulting in hypoglycosylation of substrates affecting their function and leading to disease pathologies linked to immunodeficiency (Blommaert et al., 2019; Matsuda-Lennikov et al., 2019).

Utilisation of potential glycosylation sites or sequons is not guaranteed and is dependent on the position within the chain (Nilsson and von Heijne, 2000; Ruiz-Canada et al., 2009; Shrimal et al., 2013), or the amino acid context of the site (Shrimal and Gilmore, 2013) with the kinetics of the folding or collapse of the polypeptide chain affecting glycosylation. Sequons buried within a protein structure, present at the amino or carboxy terminus or close to cysteines involved in disulfide formation may be underutilised giving rise to heterogeneity in glycoprotein forms.

Hypoglycosylation of sequons due to disulfide formation can be dependent upon STT3A or STT3B and is reversed when proteins are prevented from forming disulfides under highly reducing conditions (Allen et al., 1995; Cherepanova et al., 2014). In addition, STT3B-dependent glycosylation of cysteine-proximal sites requires the oxidoreductase activity of the thioredoxin-domain containing subunits MagT1 or TUSC3 (Cherepanova and Gilmore, 2016; Cherepanova et al., 2014). Structural analysis of TUSC3 indicates its direct binding to cysteine-containing peptides, suggesting direct binding to the polypeptide to slow down protein folding and disulfide formation to allow glycosylation to occur (Mohorko et al., 2014). The fact that MagT1 is mainly oxidised in cells (Cherepanova et al., 2014) would suggest that it acts as a reductase, thereby preventing disulfide formation prior to glycosylation. Taken together these observations indicate a crucial role for the STT3B complex in coupling disulfide formation and glycosylation to maximise utilisation of cysteine-proximal acceptor sites.

The temporal relationship between disulfide formation and glycosylation suggests that the redox status of the ER may contribute towards sequon utilisation (Cherepanova et al., 2016). ER redox reactions are balanced to allow both disulfide formation and reduction resulting in the formation of the correct disulfides within folding proteins (Bulleid and van Lith, 2014). Members of the protein disulfide isomerase (PDI) family are thioredoxin-domain containing proteins that catalyse disulfide exchange reactions (Bulleid, 2012). Their oxidation is catalysed by Ero1, which couples the reduction of oxygen to the formation of a disulfide in PDI (Cabibbo et al., 2000). Specific members of the PDI family, such as ERp57 (Jessop et al., 2007) and ERdj5 (Oka et al., 2013; Ushioda et al., 2008) catalyse the reduction of non-native disulfides either allowing the correct disulfides to form or targeting misfolded proteins for degradation. Exactly how these PDI enzymes are reduced is unknown but recent evidence suggests a role for the cytosolic reductive pathway in correct disulfide formation, driven by the reduction of thioredoxin reductase (Cao et al., 2020; Poet et al., 2017).

It is likely that the ER oxidative and reductive pathways influence the STT3B subunits, MagT1 and TUSC3 during oxidoreductase activity towards cysteines proximal to sequons. Hence, the correct utilisation of sequons may well be regulated by the prevailing redox conditions within the ER. To address the role of ER redox conditions on utilisation of STT3B-dependent acceptor sites we capitalised on a recently described *in vitro* translation system that reproduces the early stages of secretory protein ER translocation and modification under defined redox conditions (Poet et al., 2017; Robinson and Bulleid, 2020). In this system, the redox conditions can be manipulated simply by the addition of glucose 6-phosphate (G6-P) which recycles NADPH thereby driving the cytosolic reductive pathway. When a source of ER is included during translation, the newly synthesised proteins are translocated across the ER membrane and can undergo both disulfide formation and N-linked glycosylation (Wilson et al., 1995). We chose to translate the STT3B-dependent substrate hemopexin (Cherepanova et al., 2014) in such a system, and show that it is hypoglycosylated in the absence of added G6-P, an effect that is reversed upon G6-P inclusion. Our results highlight the role of ER redox in the efficiency of sequon glycosylation and reveals an unexpected role for the NADPH-dependent cytosolic reductive pathway in the function of both the STTA and STT3B-containing OST complex.

## Results

### Cytosolic reductive pathway determines the extent of sequon usage in a STT3B-dependent glycoprotein

Our initial experiments aimed to determine whether the redox conditions within the ER had any effect on the fidelity of sequon usage within a model protein, hemopexin, that had previous been shown to undergo STT3B-dependent hypoglycosylation. Hemopexin has 5 potential sequons and forms six disulfides (Fig. 1A). For these experiments we adjusted the redox status of our *in vitro* translation reactions by adding specific components to the rabbit reticulocyte lysate. We have previously shown that a commercial reticulocyte lysate that has no added DTT allows disulfides to form in proteins synthesised even in the absence of semi-permeabilised (SP) cells as a source of ER (Poet et al., 2017). Supplementing this lysate with G6-P to drive G6-P dehydrogenase (G6PDH) and thioredoxin reductase (TrxR1) activity, renders this lysate sufficiently reducing to prevent disulfide formation in proteins synthesised without SP cells but allows disulfide formation in translocated proteins when SP-cells are present. When hemopexin was translated in the absence of added G6-P and presence of SP-cells we noted the appearance of two potential glycoforms giving rise to a doublet after SDS-PAGE (Fig. 1B, lane 1). We also see an additional product of approximately 55 kDa (indicated with an arrow) that most likely corresponds to untranslocated and, therefore, unglycosylated protein. When G6-P was added to the translation, the slower migrating glycoforms predominates (lane 2). Likewise, only the slower migrating glycoform was synthesised when translations were carried out in the presence of the membrane permeable and impermeable reducing agent DTT or TCEP respectively (lanes 3 and 4). Hence it would appear that hemopexin is hypoglycosylated in the absence of G6-P, an effect that is reversed when translations were carried out under more reducing conditions. As G6-P most likely alters the redox conditions by recycling NADP to NADPH in the cytosol and TCEP is membrane impermeable these results suggest that the redox conditions on the cytosolic side of the ER membrane are affecting the glycosylation efficiency of ER translocated hemopexin.

**Figure 1.**
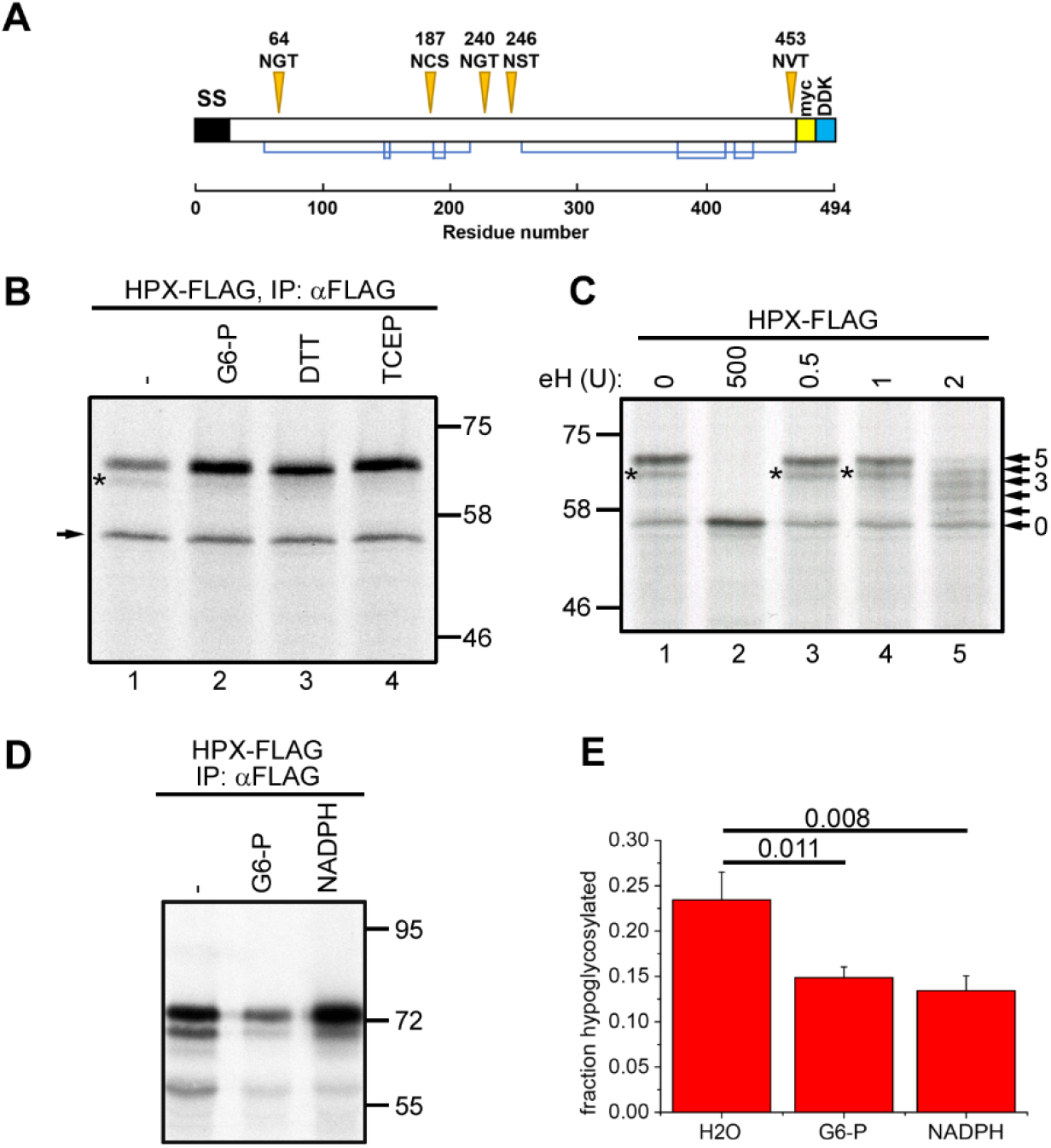
Hemopexin hypoglycosylation is prevented G6-P or NADPH in included during translation. (A) Schematic diagram of hemopexin showing glycosylation sites (orange triangles), disulfide bond connectivity (blue lines). The positions of the signal peptide (SS) and the myc- and DDK-tags are also indicated. (B) Hemopexin was translated in vitro in the presence of SP cells in the absence or presence of 5 mM G6-P, 5 mM DTT or 1 mM TCEP for 1 hour. The SP cells were washed, lysed and hemopexin was immuno-isolated with an anti-FLAG antibody, followed by separation by SDS-PAGE. The asterisk indicates hypoglycosylated hemopexin, the arrow indicates non-translocated, non-glycosylated hemopexin. (C) Hemopexin was translated as in (B) in the absence of G6-P. After lysis, the samples were incubated with different amounts of endo H (indicated as units) at 37°C for 15 minutes and analysed by SDS-PAGE. The asterisk indicates hypoglycosylated hemopexin, the arrows point to the different glycosylated species. (D) Hemopexin was translated as in (B) in the absence or presence of 5 mM G6-P or 1 mM NADPH for 30 minutes. The translation reactions were analysed as in (B). (E) Quantification of hemopexin hypoglycosylation of three independent experiments as in (D). Error bars are standard deviation of mean. p-values of student’s t-test are shown above the bar diagram.

To verify that the two translation products seen after synthesis of hemopexin are indeed glycoproteins and to identify the status of the faster migrating band, we carried out a limited digestion of the protein with endoglycosidase (endo) H (Fig. 1C). Digestion of the translation products with the highest enzyme concentration resulted in a single band corresponding to the fully deglycosylated protein indicating that the two products are indeed glycoforms (lanes 1 and 2). Addition of limiting amounts of endo H to the reaction allowed partial digestion revealing all 5 potential glycoforms that arise from variable digestion of the five oligosaccharide side chains on hemopexin (lane 5). From this analysis we can conclude that the two apparent glycoforms seen when hemopexin is translated in the absence of added G6-P are indeed the 5 and 4 glycan forms. These results are consistent with our previously observed hypoglycosylation of hemopexin when expressed in mammalian cells (Shrimal and Gilmore, 2013).

To verify that the reversal of hemopexin hypoglycosylation by G6-P is mediated by the recycling of NADP, we supplemented the translation reactions with NADPH (Fig. 1D, E). As with G6-P, we could reverse the hypoglycosylation of hemopexin just by adding NADPH confirming that the effect is not due to G6-P directly influencing the glycosylation machinery or synthesis of the oligosaccharide side chain.

### Defining the sequon giving rise to G6-P-dependent hypoglycosylation

To determine which sequon within hemopexin is hypoglycosylated, we mutated hemopexin N187 and N453 to glutamine, as it has been noted before that these sequons can frequently be skipped by the oligosaccharyl transferases (Shrimal and Gilmore, 2013). We found that hypoglycosylation in the absence of G6-P occurred with wild type hemopexin and hemopexin N453Q, which can be resolved by the addition of G6-P (Fig. 2A, lanes 1, 2 and 5, 6). No hypoglycosylation was observed with hemopexin when N187 is mutated in either the single or double mutant (lanes 3 and 7).

**Figure 2.**
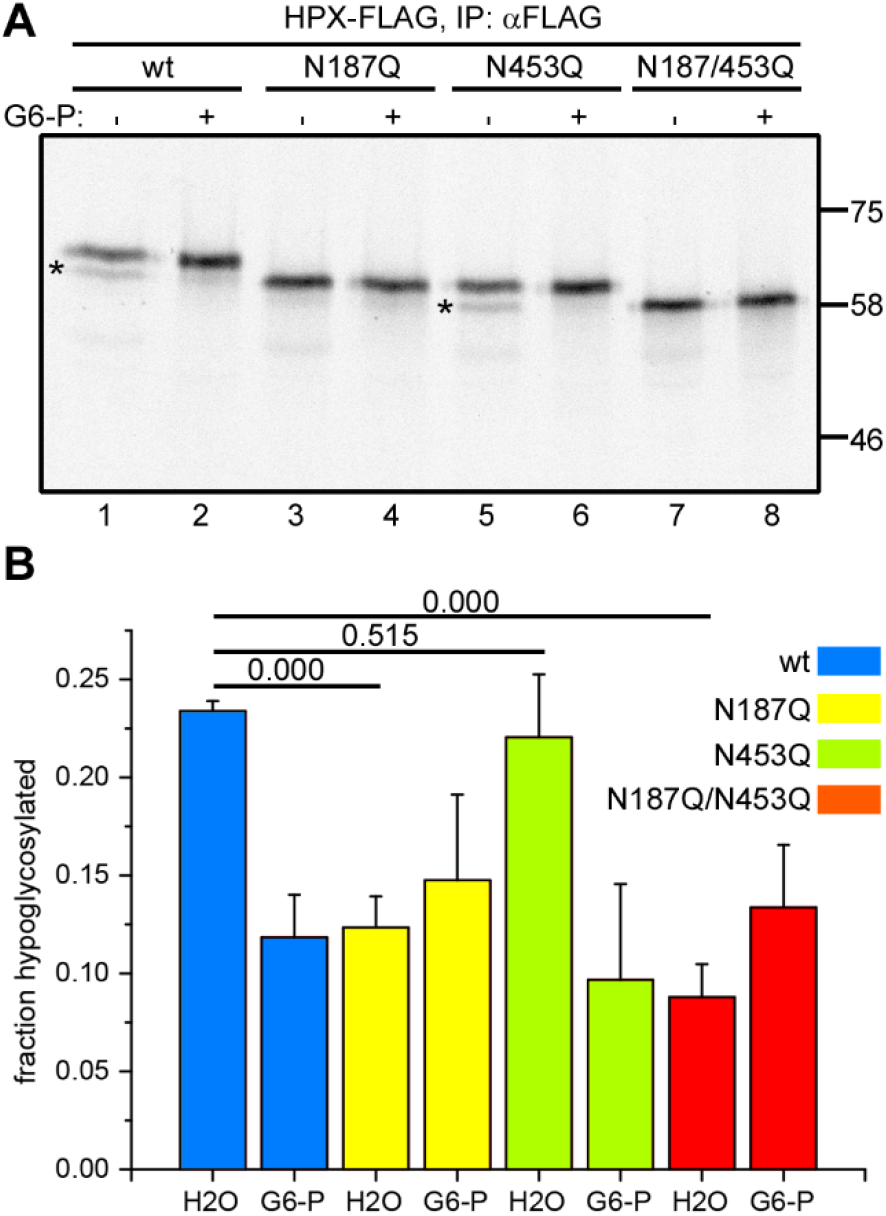
Hemopexin hypoglycosylation occurs at sequon N187. (A) Wild-type and N-glycosylation mutants of hemopexin were in vitro translated in the presence of SP cells in the absence or presence of 5 mM G6-P. The SP cells were washed, lysed and hemopexin was immuno-isolated followed by SDS-PAGE and autoradiography. Hypoglycosylated hemopexin is indicated by asterisks. (B) Quantification of hemopexin hypoglycosylation of three independent experiments as in (A). Error bars are standard deviation of mean. p-values of student’s t-test are shown above the bar diagram.

Quantification of the level of hypoglycosylation from three separate experiments supports the qualitative gel analysis (Fig. 2B). These results demonstrate that N187 is the acceptor site that is inefficiently glycosylated when translated in the presence of SP-cells and in the absence of added G6-P.

The N187 sequon is NCS with the C188 forming a short-range disulfide with C200 in the native structure of hemopexin (Fig. 1A) (Paoli et al., 1999). As it has been observed previously that cysteine-proximal glycosylation sites are often skipped (Cherepanova et al., 2014), we evaluated the role of hemopexin C188 in hypoglycosylation. In addition, to determine if the formation of the C188-C200 disulfide prevents efficient glycosylation, we mutated both cysteines individually and together to serine. We found that mutation of the more distal C200 did not prevent hypoglycosylation, which was resolved by inclusion of G6-P (Fig. 3A, lanes 5 and 6). In contrast, mutation of C188, in either C188S or C188S/C200S mutant, resulted in almost complete loss of hypoglycosylation (lanes 3 and 7).

**Figure 3.**
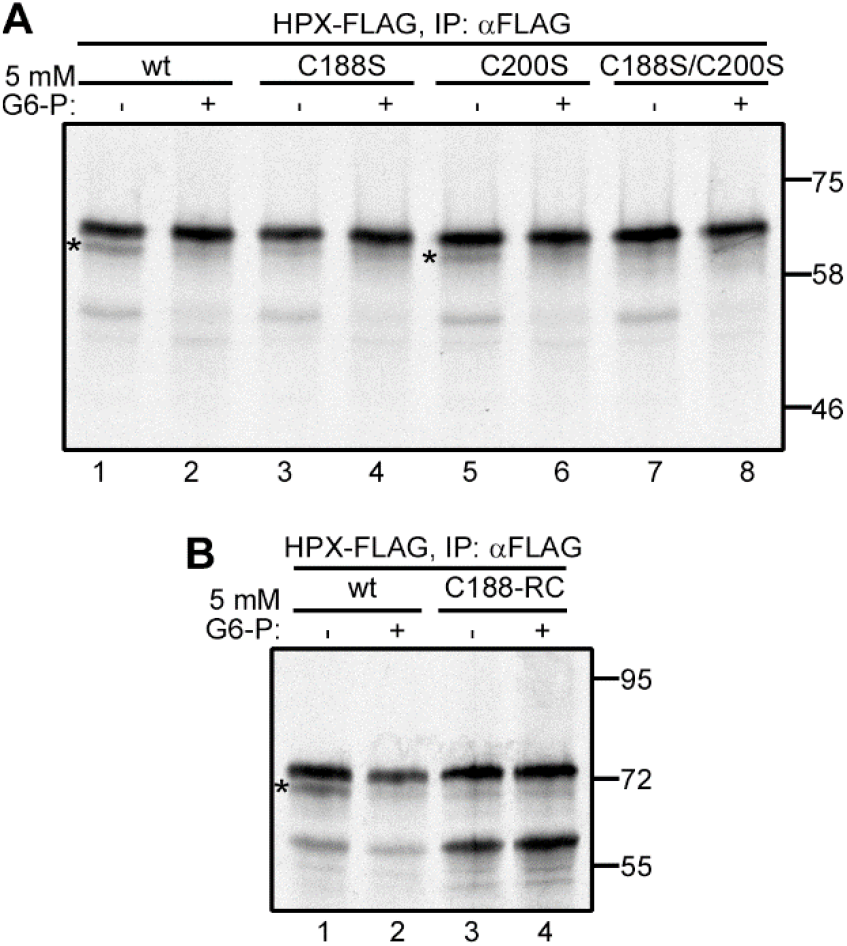
Disulfide formation via C188 results in hemopexin hypoglycosylation. (A) Wild-type and cysteine mutants of hemopexin were translated in vitro in the presence of SP cells in the absence or presence of 5 mM G6-P for 1 hour. The SP cells were washed, lysed, immune-isolated and analysed by SDS-PAGE and autoradiography. Hypoglycosylated hemopexin is indicated by asterisks. (B) Wild-type hemopexin and hemopexin lacking all mature protein cysteines except C188 were translated in vitro into SP cells in the absence or presence of 5 mM G6-P for 1 hour, and analysed as in (A).

Preventing the native disulfide from forming by mutating C200 did not stop hypoglycosylation of the acceptor site whereas mutating C188 did, suggesting that the presence of the cysteine restricts glycosylation rather than the disulfide per se. Alternatively, C188 could be oxidised or form a non-native disulfide to an alternate cysteine to C200. To test these two possibilities, we created a construct where we mutated all the cysteines in the sequence apart from the cysteine at C188. Upon translation the protein was fully glycosylated either in the presence or absence of G6-P (Fig. 3B, lanes 3 and 4). This result suggests that the formation of either a native or non-native disulfide via C188 restricts the ability of the OST from glycosylating N187 resulting in hypoglycosylation. In addition, it shows that it is a change in the redox conditions during synthesis that is reversed by the inclusion of G6-P maintaining the C188 in a reduced state to allow efficient glycosylation.

### Hemopexin forms distinct disulfide-bonded species during translation in the absence or presence of added G6-P

The ability of C188 to form a native or non-native disulfide affected N187 occupancy, which suggests a role for G6-P in modulating disulfide formation in our translation system. Indeed, we have previously shown that addition of G6-P prevents non-native disulfide formation in a variety of proteins by recycling NADPH and maintaining cytosolic thioredoxin in a reduced state (Poet et al., 2017). To determine the redox status of hemopexin following translation, we prevented disulfide rearrangement following synthesis using an alkylating agent and separated the translation products under non-reducing conditions. Typically, long-range disulfides formed in proteins affect their electrophoretic mobility by altering the hydrodynamic volume of the denatured protein. When the hemopexin translation products were analysed this way, we observed several oxidised species with a greater mobility than the reduced protein (Fig. 4A, compare lane 1 and 5). To rule out any contribution of hypoglycosylation to the pattern under non-reducing conditions, the samples were also treated with endo H to remove all glycans (even numbered lanes). Multiple oxidised species were still observed indicating that hemopexin forms distinct and incompletely disulfide-bonded species in our translation system.

**Figure 4.**
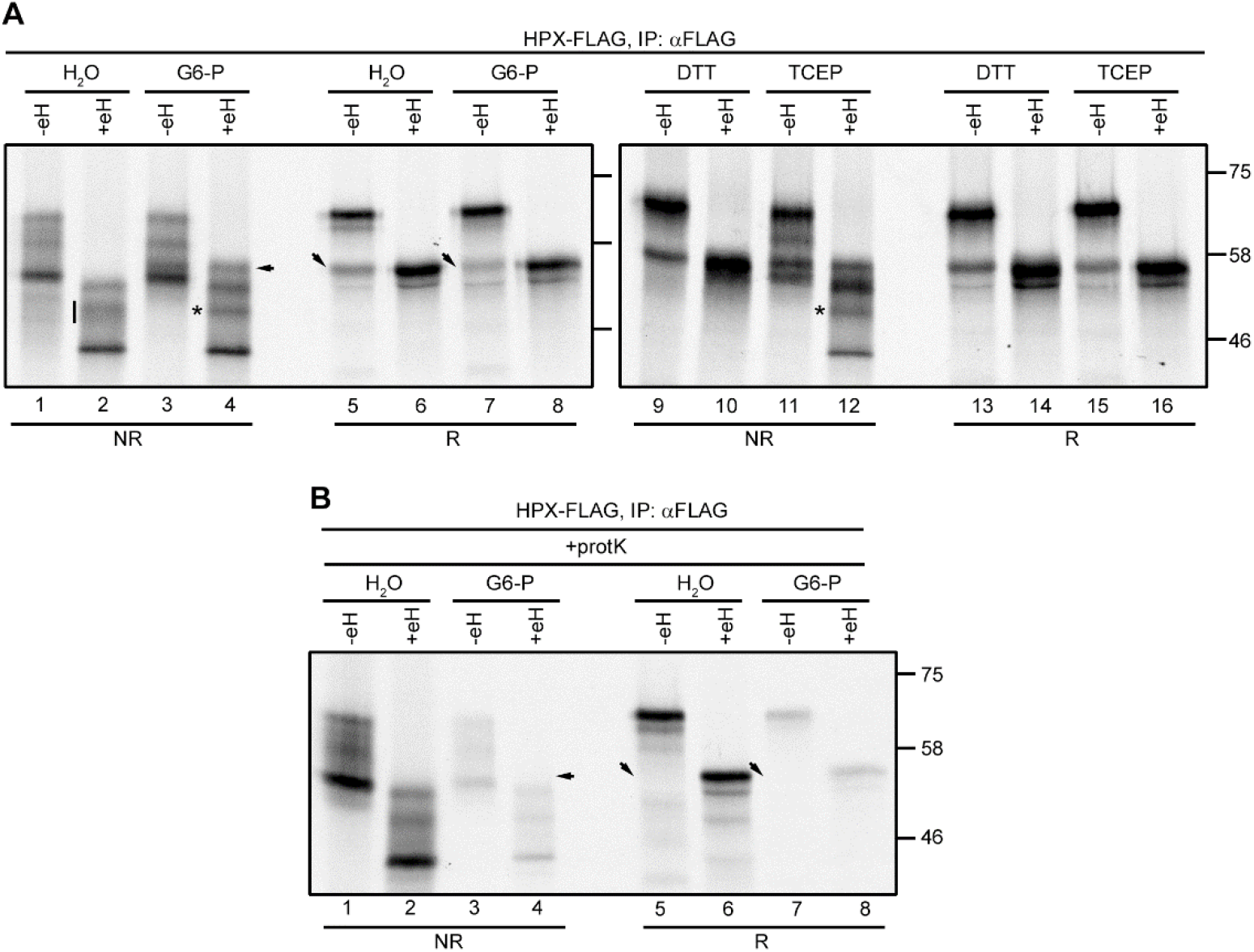
Hemopexin forms several disulfide-bonded species when translated in the presence or absence of G6-P. (A) Hemopexin was translated in vitro in the presence of SP cells in the absence or presence of 5 mM G6-P, 5 mM DTT or 1 mM TCEP. The translations were stopped by adding 20 mM N-ethyl maleimide and incubation on ice. After lysis, the samples were mock treated or treated with endo H, followed by immuno-isolation and analysis by reducing (R) and non-reducing (NR) SDS-PAGE. The vertical line and asterisks indicate a change in oxidation state when in vitro translations were carried in the presence of G6-P or TCEP. (B) Hemopexin was translated as in (A). After the addition of 20 mM NEM and pelleting the SP cells, the cells were treated with proteinase K, followed by lysis and endo H treatment. The samples were immune-isolated and analysed by reducing (R) and non-reducing (NR) SDS-PAGE. The arrows in (A) and (B) point to bands that disappear upon proteinase K treatment.

When the translations were carried out in the presence of G6-P, most of the oxidised species migrated as those in untreated lysates (lane 1 and 2 vs 3 and 4). One species appeared after G6-P addition, seen in the endoH treated samples (lane 2 versus 4, indicated by an arrow). This translation product was digested by proteinase K, indicating that it corresponds to untranslocated hemopexin (Fig. 4B, lane 4, arrow). The removal of untranslocated material is most clearly seen when the samples are separated under reducing conditions (lane 5 and 7, down arrows). The remainder of the bands are protected from proteinase K digestion, indicated that all these species are translocated into the ER lumen. The dramatic change to the redox status of untranslocated protein is consistent with our previous studies indicating that the addition of G6-P restores a robust reducing pathway in the cytosol, but does not prevent correct disulfide formation in proteins translocated across the ER membrane (Poet et al., 2017; Robinson and Bulleid, 2020). Interestingly, the oxidised species migrating with intermediate mobility become less diffuse after G6-P addition (lanes 2 and 4, vertical line and asterisk) suggesting some rearrangement of disulfides.

Adding the reductant DTT to the hemopexin *in vitro* translations prevented disulfide formation resulting in a non-reducing pattern that resembled that of the samples run under reducing conditions (Fig. 4A, lanes 9 and 10 vs 13 and 14). In contrast, the addition of the membrane impermeable reductant TCEP to the reactions gave a non-reducing pattern like that following G6-P addition (lanes 3 and 4 versus 11 and 12), including the sharpening of the intermediate oxidised species (lane 12, asterisk). Both the addition of TCEP and G6-P altered the overall pattern of translocated disulfide-bonded forms and TCEP and to some extent G6-P addition resulted in a shift in their ratios with more of the slower migrating form present following their addition (compare lanes 2, 4 and 12). Hence, G6-P inclusion caused subtle but distinct differences in the oxidised species present indicative of rearrangement of non-native disulfides. While the addition of G6-P, TCEP or DTT had different effects on the redox species formed, they all resulted in full hemopexin glycosylation.

### G6-P does not act via an ER NADPH pool or PDIs involved in non-native disulfide reduction

The results presented so far would suggest a requirement to maintain a robust cytosolic reductive pathway to ensure the efficient glycosylation of hemopexin. Alternatively, G6-P can be transported into the lumen of the ER by the glucose-6-phosphate transporter (G6PT), where it can be used by the ER-localised hexose-6-phosphate dehydrogenase (H6PDH) to locally generate NADPH (Clarke and Mason, 2003). The NADPH could be used to provide reducing equivalents for the promotion of hemopexin glycosylation by a yet unidentified pathway. To determine if the ER NADPH pool is involved, we evaluated the effect of G6-P on hemopexin glycosylation using SP-cells derived from a H6PDH knock out (KO) cell-line (Fig. 5A). G6-P addition still reversed the hypoglycosylation of hemopexin in this cell line ruling out a role for H6PDH (Fig. 5B).

**Figure 5.**
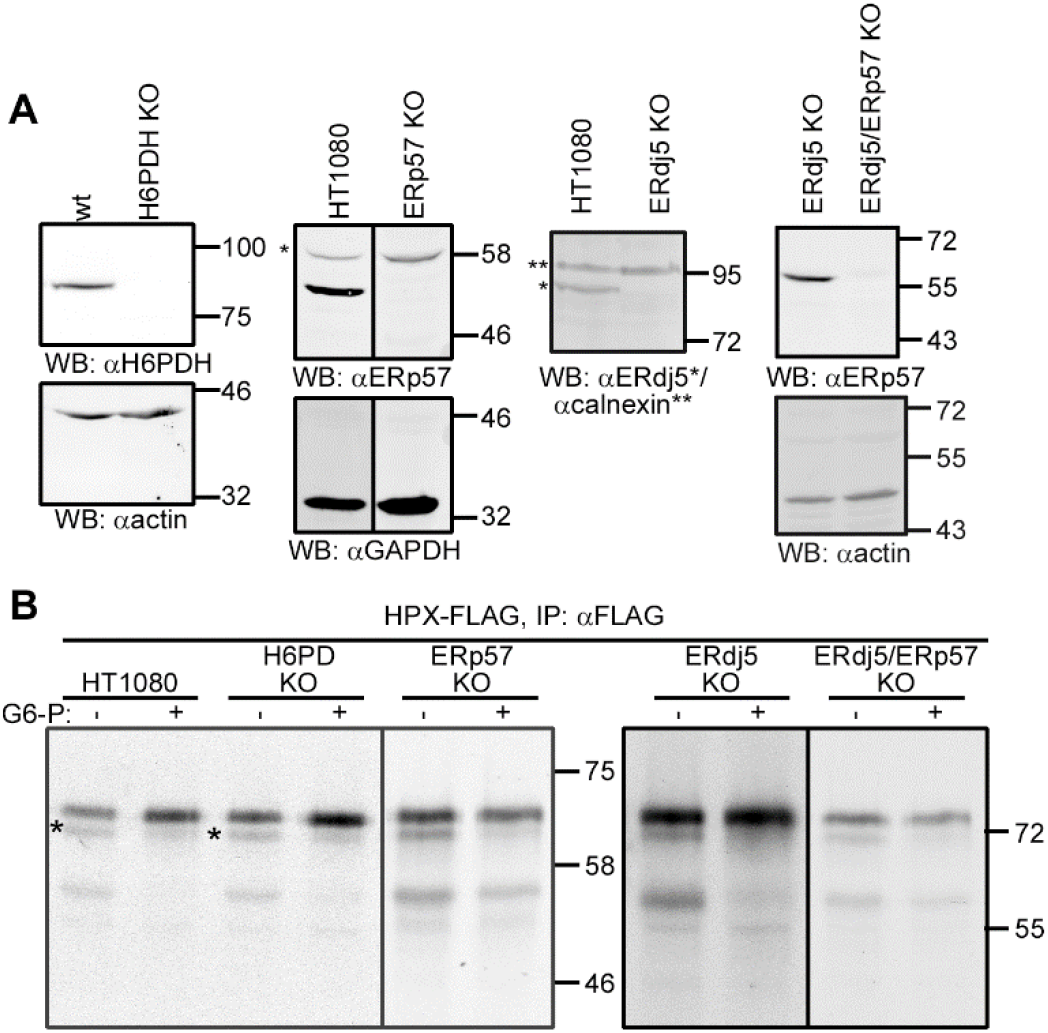
ERp57, ERdj5 or H6PDH are not required for efficient hemopexin glycosylation. (A) Western blot analysis for H6PDH, ERp57, ERdj5 and ERp57+ERdj5 knockout cell lines showing lack of expression for each knocked-out protein. Actin, GAPDH and calnexin are used as loading controls. (B) Hemopexin was in vitro translated in the presence of SP cells derived from H6PDH, ERp57, ERdj5 or ERp57+ERdj5 knockout cell lines in the absence or presence of 5 mM G6-P for 1 hour. The in vitro translations were analysed as in Fig. 3.

The fact that hypoglycosylation of hemopexin can be influenced by a disulfide formed via C188 led us to determine whether previously characterised PDIs that have reductase activity might be involved directly or via disulfide exchange with the STT3B-subunits MagT1 or TUSC3. We focused on the PDIs ERp57 and ERdj5 as they are known to catalyse the reduction of non-native disulfides (Jessop et al., 2007; Oka et al., 2013; Ushioda et al., 2008). We created KO cell-lines for the individual proteins as well as a combined ERdj5-ERp57 double KO (Fig. 5A). For each of these cell-lines there was no effect on the reversal of hypoglycosylation facilitated by G6-P (Fig. 5B). Hence the reductive pathway maintained by the addition of G6-P functions independently from the known ER PDI reductases.

### Role of the OST complexes containing either STT3A or STT3B catalytic subunits

To determine whether STT3A or STT3B are required for the G6-P effect on hemopexin glycosylation, we used two previously characterised KO cell lines for STT3A and STT3B (Cherepanova and Gilmore, 2016). To look specifically at glycosylation of the cysteine proximal N187 we used the hemopexin N453Q mutant for *in vitro* translation in SP cells derived from these KO cell lines.

Previously, the N453 site had been shown to be STT3B-dependent (Shrimal and Gilmore, 2013). Hemopexin N453Q translated into STT3A KO SP cells was glycosylated similarly to that in wild type cells, with both G6-P and DTT able to partially resolve the hypoglycosylation (Fig. 6A, lanes 4-6 versus 1-3 and Fig. 6C). These results indicate that the effect of G6-P seen in wild type cells is not due to the STT3A OST.

**Figure 6.**
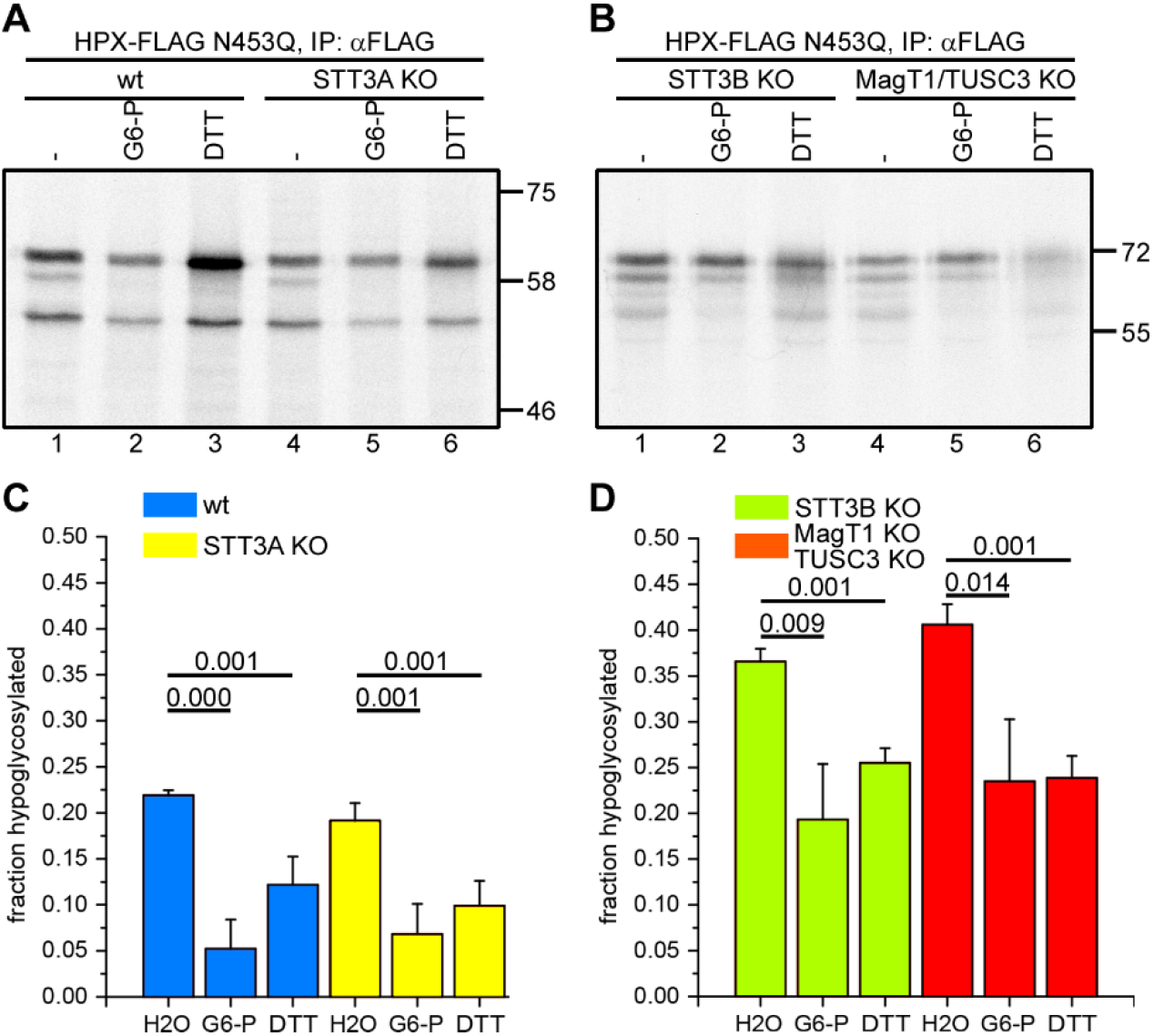
STT3A and STT3B dependence on hemopexin glycosylation. (A)(B) Hemopexin N453Q was translated in vitro in the presence of SP cells derived from wild-type, STT3A, STT3B or MagT1/TUSC3 KO cells in the absence or presence of 5 mM G6-P or 5 mM DTT at 30°C for 1 hour. After immuno-isolation, hemopexin was analysed by SDS-PAGE. (C)(D) Quantification of hypoglycosylated hemopexin of 4 and 3 independent experiments for (A) and (B), respectively. Error bars are standard deviation of mean. p-values of student’s t-test are shown above the bar diagram.

In contrast, hemopexin hypoglycosylation is more pronounced in STT3B KO cells in the absence of G6-P with levels of 35-40% hypoglycosylation as compared with 20-25% in wild type cells (Fig. 6B and D compared to Fig. 6A). There was an effect of adding G6-P or DTT which reduced the hypoglycosylation to wild-type levels seen in the absence of reducing agent (Fig. 6D). In STT3B knockout cells, not only is the STT3B catalytic activity abolished, but these cells have also lost expression of the STT3B-specific subunits MagT1/TUSC3 (Cherepanova and Gilmore, 2016). To see if there is a direct contribution of MagT1/TUSC3, we used SP cells derived from MagT1/TUSC3 KO cells for *in vitro* translation of hemopexin. Like with the STT3B KO cells, hemopexin hypoglycosylation was more pronounced in the absence of G6-P and, when either G6-P or DTT was included during translation, hypoglycosylation was reduced to the levels seen in wild type cells in the absence of G6-P (Fig. 6B, lanes 4-6 and Fig. 6D). Thus, efficient glycosylation of hemopexin N187 is dependent on the STT3B complex and specifically the TUSC3 or MagT1 subunits.

## Discussion

Previous work demonstrated a role for the STT3B-associated oxidoreductase subunits MagT1 or TUSC3 in ensuring the efficient glycosylation of the STT3B-dependent substrate hemopexin (Cherepanova et al., 2014). Here we extend this work to demonstrate that the redox conditions within the ER contribute towards sequon utilisation and that the cytosolic reductive pathway plays a key role in maintaining the optimal redox balance for STT3B activity. We find that the efficiency of cysteine-proximal acceptor site usage is likely dependent upon the timing of disulfide formation. If a disulfide forms prior to glycosylation by the STT3A containing OST, the site may be skipped and requires a STT3B dependent reduction of the disulfide to allow glycosylation. In addition, the function of STT3B containing OST is dependent upon the thioredoxin-like subunits MagT1 or TUSC3 and their activity in reducing any disulfides requires a robust reductive pathway. Hence, the cytosolic reductive pathway influences the initial glycosylation by STT3A by delaying disulfide formation as well as optimising STT3B function in glycosylation of sites missed by STT3A due to rapid disulfide formation. The cytosolic reductive pathway is itself dependent upon the presence of an active pentose phosphate pathway to ensure the recycling of NADPH illustrating a connection between cellular metabolism and glycosylation efficiency. While such a link has been suggested previously (Gansemer et al., 2020), our observations provide a previously unappreciated correlation between cellular metabolism and the variability in synthesis of protein glycoforms.

If the ability of STT3A to glycosylate the N187 sequon is determined by whether a disulfide has formed, this would indicate that the STT3B containing OST needs to first reduce any disulfide between C188 and C200 prior to glycosylating this site. Hence, the MagT1 or TUSC3 subunits would act as reductases which is suggested by the relatively low reduction potential of its active site thiols and by the fact that it is predominantly in an oxidised form in the ER (Cherepanova et al., 2014; Mohorko et al., 2014). In addition, substrate trapping mutants of MagT1, where the second cysteine in the CXXC active site was mutated to serine, form mixed disulfides to substrate proteins, further confirming the ability to act as a reductase (Cherepanova et al., 2014). For the enzyme to function as a reductase its active site must be reduced either by a low molecular weight thiol such as glutathione or by disulfide exchange with another protein. The reduction potential of glutathione is higher than MagT1 (Mohorko et al., 2014) so it would be a thermodynamically unfavourable reaction, but cannot be ruled out due to the high cellular glutathione concentration (Jensen et al., 2009). Strong candidates for disulfide exchange reactions would be members of the ER-localised PDI family, though two of the previously characterised reductases ERp57 and ERdj5 do not appear to influence the G6-P dependent hypoglycosylation of hemopexin. Due to the large repertoire of PDI family members (Hatahet and Ruddock, 2007), it is likely that there is redundancy in their ability to facilitate disulfide exchange making the identification of reductases for MagT1 or TUSC3 challenging.

The transfer of reducing equivalents across the ER membrane is most likely brought about by a transmembrane protein (Cao et al., 2020), which could directly or indirectly reduce MagT1 or TUSC3. The requirement for such a conduit for reducing equivalents has been demonstrated to allow native disulfides to form in proteins entering the secretory pathway (Poet et al., 2017). Here, the presence of such a transmembrane protein is suggested by the ability of added NADPH to reverse hypoglycosylation and the fact that the ER membrane is essentially non-permeant to either NADP or NADPH (Piccirella et al., 2006). This fact, combined with the lack of a role for the ER-localised H6PDH, would suggest an indirect effect on ER redox status facilitated by the transfer of reducing equivalents across the ER membrane. In addition, the ability to reverse hypoglycosylation with the membrane impermeable reducing agent TCEP suggests an indirect effect on OST function. Whether the same membrane components are required to ensure the fidelity of disulfide formation and STT3B containing OST function remains to be determined.

## Materials and Methods

### Cell lines, constructs, and reagents

The HT1080 and 293 HEK cell lines were cultured in DMEM (Gibco) supplemented with 2 mM glutamine,100 U/ml penicillin, 100 µg/ml streptomycin and 10% foetal calf serum (Sigma). The wild-type, STT3A, STT3B and MagT1/TUSC3 293 HEK knockout cell lines were created as described (Cherepanova and Gilmore, 2016). The antibodies against FLAG-tag (Sigma F3165), H6PDH (Abcam ab119046), actin (Sigma A2103), DNAJC10 (ERdj5) (Abnova, H00054431-M01A) and GAPDH (Ambion AM4300) were obtained commercially. The antibody to ERp57 was raised as described (Jessop and Bulleid, 2004). Myc- and DDK-tagged hemopexin plasmid was purchased from Origene. Glycosylation and cysteine mutations were made with the Quick Change II Site-Directed Mutagenesis kit (Promega). A synthetic construct coding for the hemopexin sequence without any cysteines apart from cys188 was purchased from Genescript. Glucose-6-phosphate, N-ethylmaleimide and DTT were obtained from Sigma. Tris(2-carboxyethyl) phosphine (TCEP) was purchased from Thermo Fisher.

### *In vitro* transcription and *in vitro* translation

Myc- and DDK (FLAG)-tagged Hemopexin mRNA was transcribed from an AgeI (NEB) linearised plasmid using T7 RNA polymerase (Promega). *In vitro* translation (Poet et al., 2017) and preparation of semi-permeabilised (SP) cells (Wilson et al., 1995) was carried out as described previously. Briefly, the mRNA was translated in a Flexi rabbit reticulocyte lysate (Promega) containing ^35^S methionine/cysteine (Perkin Elmer) in the presence of SP-cells at 30°C for 1 hour. Where indicated the translation reaction was supplemented with G6-P (5 mM), DTT (5 mM), TCEP (1 mM) or NADPH (1 mM). The SP cells were pelleted by centrifugation and washed with KHM buffer (110 mM KAc, 2 mM MgAc, 20 mM Hepes pH 7.2) followed by lysis, preclearing with agarose beads and immunoprecipitation with anti-FLAG antibody and protein G beads (Generon). The samples were analysed by SDS-PAGE and visualised by exposure to film (Kodak) or phosphorimager plates (Fujifilm FLA-7000). Quantifications were performed with ImageJ (https://imagej.nih.gov/ij/) using phosphorimager scans and were calculated as the fraction that is hypoglycosylated (hypo/(hypo+full)).

To distinguish between translocated and non-translocated translation products, after translations the SP cells were incubated on ice with 10 µg/ml proteinase K (Roche) in the presence of 1 mM CaCl_2_ for 25 minutes. The digestion was stopped with 0.5 mM PMSF before further analysis. Where indicated, the lysates were treated with endoglycosidase H (NEB).

### CRISPR-Cas 9 knockout cell-lines

Hexose-6-phosphate dehydrogenase (H6PDH) knockout cells were created by CRISPR-Cas9, using a two-guide (GGATTATGGAGACATGTCCC and GCCATAAGTACTTCTTAGCC) Cas9 D10A nickase system (Kabadi et al., 2014). The same system was used for the creation of the ERp57 KO cells using the guide sequences (TTCTAGCACGTCGGAGGCAG and CGAGAGTCGCATCTCCGACA). The ERdj5 knockout cells were created using the Integrated DNA Technologies (IDT) Alt-R CRISPR-Cas 9 System, the predesigned gRNA to knock out ERdj5 ([GTGTATATGGCCATTTTAGT](https://www.ensembl.org/Homo_sapiens/ZMenu/TextSequence?db=core;factorytype=Location;g=ENSG00000077232;r=2:182716041-182794464;t=ENST00000264065;v=rs765301178;vf=205613368)) was selected. We used a single gRNA (cRNA) duplexed with a tracrRNA (IDT, 1072532) and Alt-R S.p. HiFi Cas9 Nuclease V3 (IDT, 1081060) for genome editing. To generate an ERp57 knockout, Cas9 nuclease was added to a duplex of 1AB crRNA:tracrRNA in Optimem to form the RNP complex (Fisher, 31985062), which was then transfected into HT1080 cells using Lipofectamine CRISPRMAX transfection reagent (Thermo Fisher). Cells were transferred to 15 cm dishes until colonies appeared (about 10-12 days later). All positive knockout cells were identified by western blotting using either anti-DNAJC10 (ERdj5), anti-H6PDH or anti-ERp57 and deletion verified by sequencing. The ERp57/ERdj5 knockout cell-line was created from the ERdj5 KO cells using the Alt-R CRISPR-Cas 9 system, with the ERp57 gRNA (GTCCGTGAGTTCTAGCACGT) (IDT).

## Acknowledgements

This work was funded by the Wellcome Trust (grant number 103720) and Medical Research Scotland (grant number: VAC-1417-2019).

## Author contributions

The work described in this manuscript was conceived and supervised by NJB with contribution from MvL, and RG. The experimental work was carried out by MvL, MAP, BF, GG and JI. The data was analysed by MvL and NJB. The manuscript was written by NJB and MvL with editing by RG.

## Conflict of interest

The authors declare that they have no conflict of interest.

## Notes

### Competing Interest Statement

The authors have declared no competing interest.

